# CRISPR-Cas9 interrogation of a putative fetal globin repressor in human erythroid cells

**DOI:** 10.1101/335729

**Authors:** Jennifer Chung, Wendy Magis, Jonathan Vu, Seok-Jin Heo, Kirmo Wartiovaara, Mark C. Walters, Ryo Kurita, Yukio Nakamura, Dario Boffelli, David I. K. Martin, Jacob E. Corn, Mark A. Dewitt

**Author notes:** co-corresponding authors: Mark DeWitt, Jacob Corn, David Martin. These authors contributed equally to this work (Jennifer Chung, Wendy Magis, Jacob E. Corn).

## Abstract

Sickle Cell Disease and ß-thalassemia, which are caused by defective or deficient adult ß-globin (HBB) respectively, are the most common serious genetic blood diseases in the world. Expression of the fetal ß-like globin, also known as γ-globin, can ameliorate both disorders by serving in place of the adult ß-globin. Here we use CRISPR-Cas9 gene editing to explore a putative γ-globin silencer region identified by comparison of naturally-occurring deletion mutations associated with up-regulated γ-globin. We find that deletion of a 1.7 kb consensus element or select 350 bp sub-regions from bulk populations of cells increases levels of fetal hemoglobin (HbF) or γ-globin. Screening of individual sgRNAs in one sub-region revealed three single guides that caused mild increases in γ-globin expression. However, clonal cell lines with the 1.7 kb region deleted did not up-regulate γ-globin and neither did lines with either of two of sub-regions identified in the screen deleted. These data suggest that the region is not an autonomous γ-globin silencer, and thus by itself is not a suitable therapeutic target in the ß-hemoglobinopathies.

## Introduction

The ß-hemoglobinopathies Sickle Cell Disease (SCD) and ß-thalassemia are genetic blood diseases characterized by defective or deficient adult ß-globin (*HBB*). Sickle Cell Disease (SCD) is a monogenic recessive disorder that affects at least 90,000 predominantly African-American individuals in the US *(1)*. The sickle mutation is most common in sub-Saharan Africa, where the numbers of children born with the disease are high but death in the first five years of life is common *(2, 3)*; there are also very large numbers of cases in India. The molecular basis and inheritance of SCD have been known for nearly 70 years *(4, 5)*, but even in the developed world individuals with SCD experience a greatly reduced quality of life, and suffer an ∼30-year decrement in lifespan *(1, 6)*.

ß-thalassemia is characterized by reduced or absent expression of HBB. The disease also affects hundreds of thousands worldwide, with approximately 23,000 births annually in its most severe form, transfusion-dependent ß-thalassemia *(6)*, which without chronic erythrocyte transfusions is usually fatal in the first few years of life. In ß-thalassemia, deficient ß-globin expression results in toxic free *α*-globin chains that prevent the completion of erythrocyte differentiation, resulting in profound anemia *(7)*. The disease is typically managed with chronic blood transfusions, which eventually leads to iron overload. Currently the only cure for both SCD and beta thalassemia is allogeneic stem cell transplantation *(8)*.

Both SCD and transfusion-dependent beta thalassemia can be ameliorated by increased expression of γ-globin, the ß-like globin found in fetal hemoglobin (HbF) *(10)*. In hereditary persistence of fetal hemoglobin (HPFH), mutations in the ß-globin locus cause expression of high levels of γ-globin throughout life *(11–13)*. HbF inhibits polymerization of sickle hemoglobin (HbS), and levels of HbF above ∼20% are associated with a virtual absence of manifestations of SCD. Increased HbF in adults is a positive predictor of survival in patients with SCD *(14)*. In ß-thalassemia, expression of γ-globin can replace the function of deficient ß-globin.

CRISPR-Cas9 is a recently developed gene editing technology that enables efficient genome manipulation via targeted generation of a double strand break (DSB) *(9, 10)*. The break is repaired through either of two general pathways: error-prone non-homologous end joining (NHEJ), or accurate homology-directed repair (HDR) *(11, 12)*. Targeting is directed by a 20 nucleotide guide RNA (gRNA) bound by Cas9 and a Protospacer Adjacent Motif (PAM) within the genome *(10)*. The development of this modular, user-friendly targeting system has led to an explosion of interest in gene editing, including for the treatment of the hemoglobinopathies.

Strategies to drive γ-globin expression in erythroid cells via *ex vivo* gene editing of hematopoietic stem/progenitor cells (HSPCs) have recently emerged *(13)*. Decreased expression of *BCL11A*, a transcription factor that is necessary for silencing of γ-globin expression in adult-stage erythrocytes *(14, 15)*, leads to high-level expression of γ-globin. Other efforts have targeted the ß-like globin locus, reproducing naturally occurring HPFH mutations in the promoters of the two γ-globin genes (*HBG1* and *HBG2*) *(16–18)*.

We used gene editing to explore an approach to creation of an HPFH phenotype by targeting an intergenic region upstream of the *HBD* gene. Comparison of naturally-occurring deletions causing either HPFH or ß-thalassemia identified a candidate 1.7 kb repressor region *(19)* (Figure S1). Deletion of this region and certain, smaller sub-regions in bulk pools of cells induced expression of HbF. However, multiple clonal HUDEP-2 sublines harboring a deletion of this region did not exhibit increased HbF. When the deletions were induced in hematopoietic stem and progenitor cells (HSPCs), and these cells were differentiated into erythroid colonies, we observed little up-regulation of γ-globin expression. These results suggest that the 1.7 kb region is not a stable repressor for developmental silencing of γ-globin.

## Results

We began by defining the minimal region upstream of *HBD* whose deletion is associated with increased levels of HbF. Individuals lacking the intergenic region between the δ -globin and γ-globin genes, in particular the 7.2 kb “Corfu” deletion *(20)*, and several more extended deletions including HPFH-1 and the French HPFH deletion *(17, 21)*, exhibit an HPFH phenotype. In contrast, deletions that do not include this region are not associated with high HbF expression and instead cause ß-thalassemia, as highlighted in recent work *(19)*. We identified a 1.72 kb genomic region between the 5’ breakpoint of the French HPFH deletion and the Macedonian δ/ß-thalassemia deletion using the coordinates in the HbVar online database *(19)* (Figure S1). We reasoned that CRISPR/Cas9-mediated deletion of this putative repressor region (PRR) may lead to clinically relevant levels of HbF expression (Figure 1A). Furthermore the limited size is amenable to further dissection, unlike larger deletions targeted in other studies *(17, 18, 22)*. In sum, we sought to explore the relationship between PRR deletion *genotypes* and the globin expression *phenotype*.

**Figure 1:**
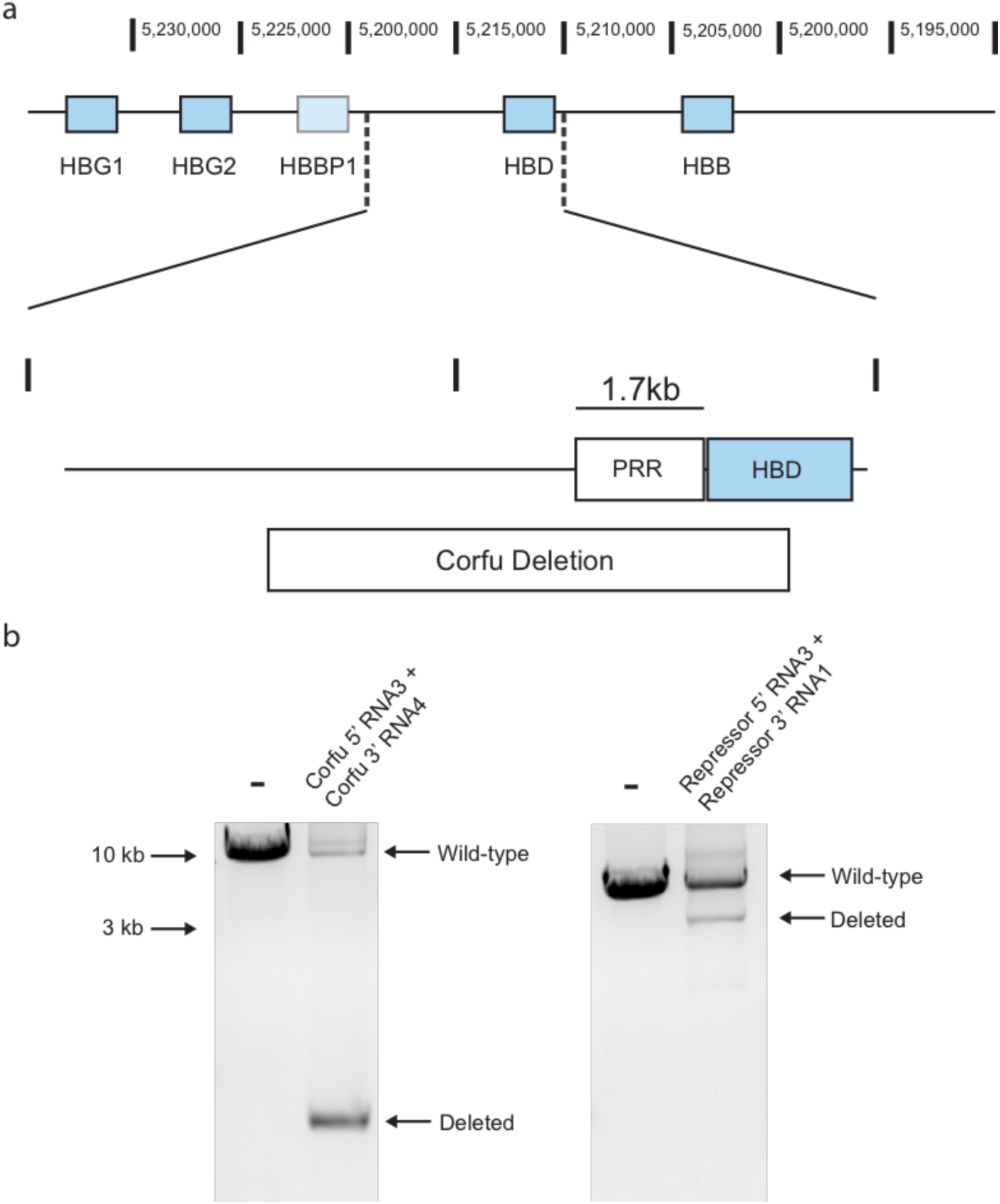
Targeted deletion of genomic regions implicated in γ-globin silencing using selection-free Cas9 RNP electroporation. A) Schematic depicting two targeted regions: the naturally-occurring Corfu HPFH deletion and the 1.7 kb putative repressor region (PRR) upstream of HBD *(13)*. B) PCR amplification after gene editing with pairs of Cas9 RNPs that delete the Corfu and PRR regions. Deletion is indicated by the presence of shorter bands corresponding to the deleted alleles.

As a model system to assess HbF expression we used the recently-developed HUDEP-2 cell line *(23)*, which readily differentiates into late-stage erythroblasts. These cells display an adult pattern of globin expression, with *HBB* expressed and *HBG* silenced, but can express *HBG* after various genetic manipulations in *cis* or *trans* to the ß-globin locus; they have been used productively to explore genotype-phenotype relationships related to globin switching *(16, 24, 25)*. To edit HUDEP-2 cells, we used Cas9 RNP electroporation, which we have found to be effective at gene targeting in CD34+ HSPCs *(26)*. Our goal was to screen the region in depth with RNP reagents to identify short regions whose deletion, perhaps with a single RNP, may activate γ-globin expression. We designed Cas9 RNPs and Cas9 RNP pairs to target progressively smaller regions, starting with the full PRR, moving to overlapping sub-regions of the PRR, and culminating in individual Cas9 RNP electroporation of a single sub-region. We edited HUDEP-2 cells, cultured them in differentiation conditions *(23, 24)*, and measured HbF expression by intracellular flow cytometry.

We generated Cas9 RNP pairs that cut at the 5’ and 3’ ends of the PRR, and of the naturally occurring Corfu deletion (Figure 1B and S2, guides in Table S3 *(20)*). Efficient cutting by individual candidate guide RNAs was assayed with T7 endonuclease I, and guides with >50% cutting at each end were paired (Figure S2). Deletion of the PRR or Corfu region in cell pools was confirmed by PCR (Figure 1B). HUDEP-2 cells that had been electroporated with RNP pairs were differentiated into erythrocytes to assess HbF expression by flow cytometry. The edited cell pools displayed an increased proportion of cells expressing HbF (Figure 2A, and Figure S3) *(27)*. 17.2% of cells expressed HbF when the PRR deletion RNPs were delivered, and 23% of cells when the Corfu deletion RNPs were delivered.

**Figure 2:**
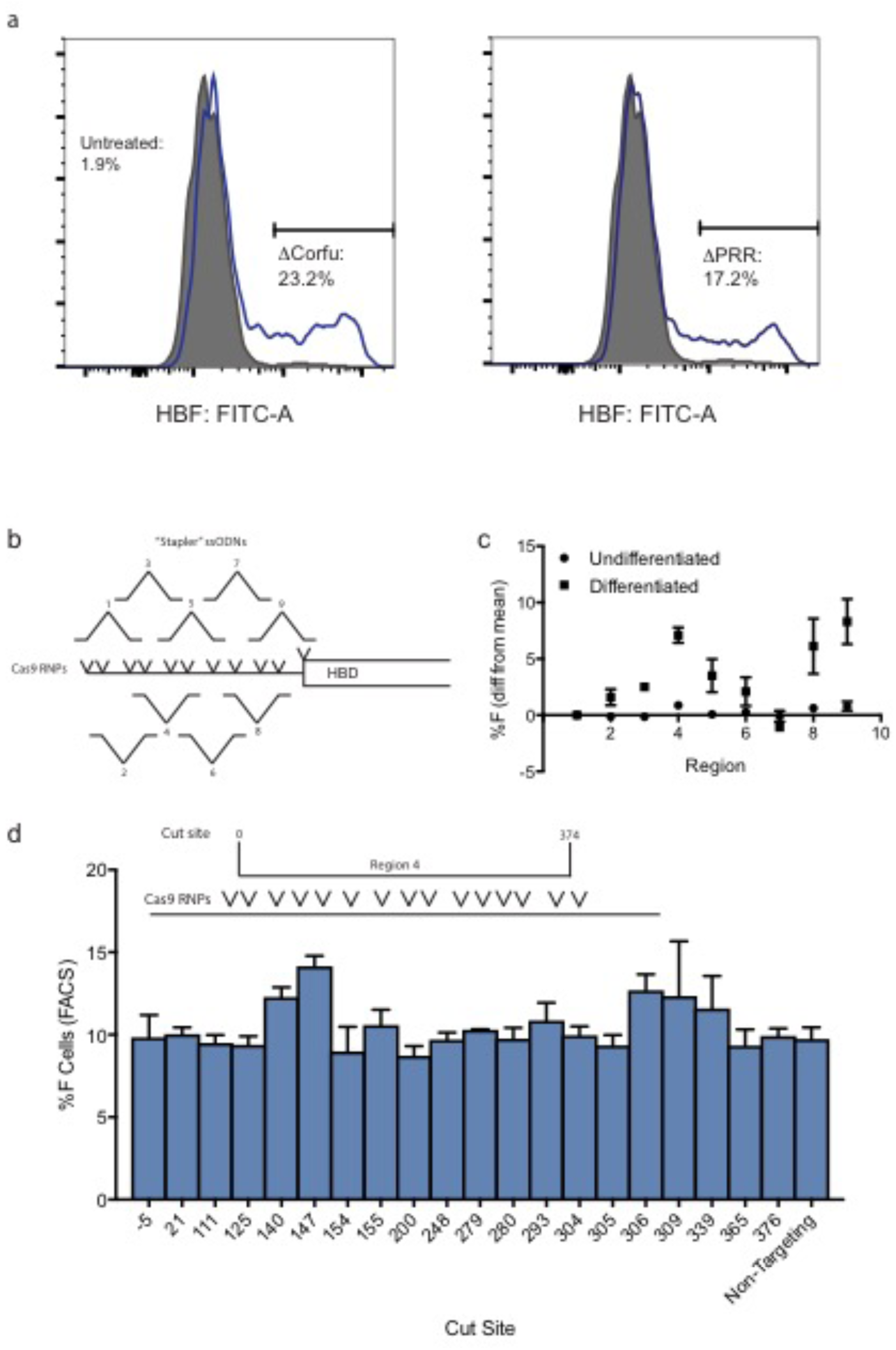
Interrogation of the PRR in the parent HUDEP-2 cell line. A) Representative intracellular FACS plots showing a population of HbF-expressing HUDEP-2 cells, after electroporation of RNP pairs generating each deletion and differentiation into erythrocytes. B) Schematic depicting the PRR, divided into 9 overlapping sub-regions. Deletion of each sub-region is programmed by a pair of RNPs and an ssDNA “stapler” containing the deletion breakpoint. C) Flow cytometric evaluation of HbF-expressing HUDEP-2 cells after introduction of Cas9 RNPs driving deletion of each sub-region, before and after differentiation into erythroblasts. Results are mean-population mean ± s.d. for 3 biological replicates. D) Scanning mutagenesis screen using 20 separate Cas9 RNPs targeting every PAM in region 4 to HUDEP-2 cells, with HbF-expressing cells enumerated by flow cytometry. To ensure consistency, synthetic RNAs were used. Results are mean ± s.d. for 3 biological replicates.

We further interrogated the PRR by dividing it into 9 overlapping regions and targeting each sub-region separately. In an attempt to enhance deletion, we designed 150 nucleotide ssODNs in which one half matched 75 bp upstream of one breakpoint, and the other half matched 75 bp downstream of the other breakpoint, thus corresponding to the sequence of a deleted allele (Table S3). We reasoned that these “stapler ssODNs” could increase the likelihood that the CRISPR-generated DSBs are resolved in favor of a precise deletion (Figure 2B). We evaluated guides by T7E1 digest of genomic DNA. Cultures of HUDEP-2 cells were treated with each pair and the appropriate stapler ssODN, and then differentiated. Of each RNP/ssODN set, those programming deletion of regions 4 and 9 were associated with a significant increase in the frequency of HbF-positive cells by flow cytometry, increasing expression by 7% above the mean (of all cultures) for region 4, and 6.5% for region 9 (Figure 2C). RNPs that delete overlapping regions 3, 5, and 8 also led to an increase. Based on these observations, we focused our subsequent studies on the region associated with the largest increase in fetal hemoglobin expression, Region 4.

Deletion of genomic elements using pairs of Cas9 RNPs is useful for defining regions of phenotypic interest. However, cutting twice in one targeted cell can lead to unwanted translocations or inversions, in addition to short indel mutations at either site, as evidenced from NGS genotyping of heterozygous edited clonal cell lines (Table S2). To avoid these complications, a single cut site is preferred. Hence we attempted to identify a single guide targeting within the PRR and capable of inducing a significant increase in HbF expression, focusing on sub-region 4. We synthesized Cas9 RNPs to target every PAM within the region (Figure 2D and Table S3). We introduced each of these 20 RNPs individually to HUDEP-2 cells by electroporation, and analyzed the effect as with the deletions discussed above (Figure 2D). Electroporation with three guides gave modest but statistically significant increases in the frequency of HbF-expressing cells, when compared to a non-targeting control RNP.

The screens presented above suggested that the full PRR, and discrete sites within it, may be necessary for silencing of γ-globin in HUDEP-2 cells. However, a given RNP pair can generate many alleles, including indels at either breakpoint, along with inversions and translocations. We therefore examined detailed genotype-phenotype relationships through generation and evaluation of clonal cell lines lacking either the PRR or sub-regions 4 and 9.

HUDEP-2 cells are a polyclonal pool of erythroblasts from a single individual, immortalized by lentiviral transduction. Clonal cell lines derived from HUDEP-2 may have subtle variations in phenotype associated with lentiviral transgene insertion, and not the CRISPR/Cas9-generted deletion of the PRR or its sub-regions *(23)*. To eliminate this as a source of variation, we made a HUDEP-2 subclone by limiting dilution, henceforth termed “H2.1”. This and the parental HUDEP-2 cell line were used in subsequent studies.

We delivered the repressor deletion RNP pairs to a HUDEP-2 pool and to H2.1, isolated edited clones by limiting dilution, and genotyped the resulting clonal cell populations by PCR amplification and multiplexed next-generation amplicon sequencing, obtaining multiple heterozygous and homozygous deletion clones from both HUDEP-2 and H2.1 (Table S1). We differentiated several sets of these clones, and assessed globin expression by flow cytometry for HbF-expressing cells and HPLC for hemoglobin expression. Surprisingly, we found that even homozygous deletion of the putative repressor did not reliably increase the percentage of HbF-expressing cell in cloal cell lines derived from both H2.1 and HUDEP-2 (Figure 3A), and failed to induce HbF at all in an HPLC assay in clones derived from H2.1 (Figure 3B). These results, which are in contrast to results in pools of edited cells, suggest that the PRR alone is not necessary for γ-globin silencing in HUDEP-2 cells.

**Figure 3:**
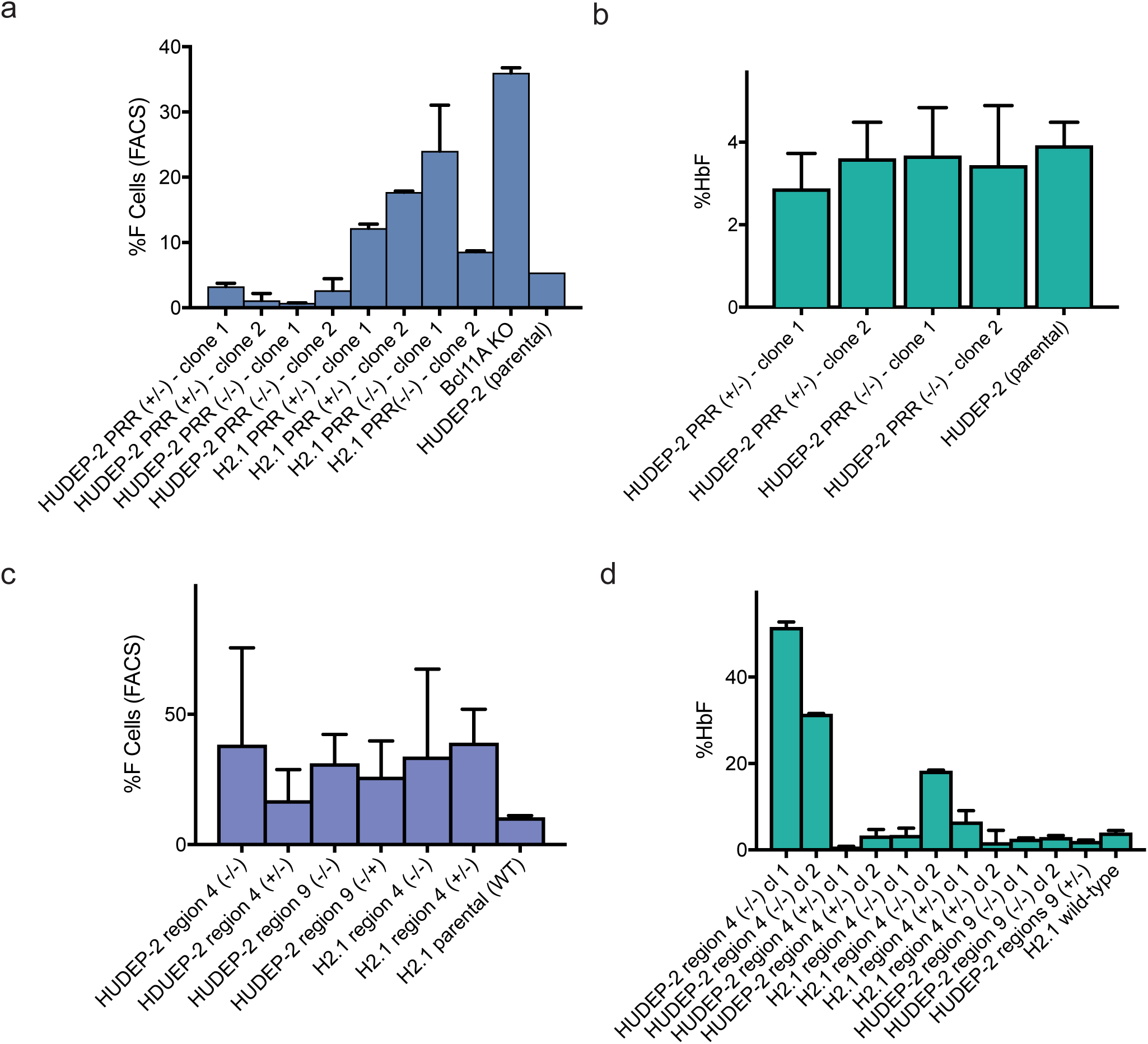
Interrogation of HUDEP-2 and H2.1 cell clones carrying deletions of the PRR and selected sub-regions. A) Proportion of HbF-expressing cells assayed by flow cytometry in cell clones carrying the indicated PRR deletions (mean ± s.d. of 3 biological replicates). B) Quantitation of HbF expression by HPLC in selected clones carrying the indicated PRR deletions, (mean ± s.d. of 3 technical replicates). No significant differences in HbF expression were seen in comparison to parent HUDEP-2 cells. C and D) Expression of HbF in clonal cell lines carrying the indicated sub-region deletions, measured by FACS (C) and HPLC (D). All clones were assayed by flow cytometry, and the average for each genotype is shown (mean ± s.d.); a subset were analyzed individually by HPLC (mean ± s.d. of 3 technical replicates).

We also generated clonal cells with deletions of the two candidate sub-regions of the PRR (4 and 9) identified by screening in the parent HUDEP-2 line (Figure S3), editing both HUDEP-2 (sub-region 4 and 9) and H2.1 (for sub-region 4 only); deletions were confirmed by next-generation sequencing (Table S1). We obtained 8 homozygous deletion and 9 heterozygous deletion clones of region 4 in HUDEP-2, 2 homozygous and 7 heterozygous deletion clones of region 4 in H2.1, and 3 homozygous and 2 heterozygous deletion clones of region 9 in HUDEP-2. These clones were differentiated, and HbF-expressing cells were counted by flow cytometry and total HbF expression was measured with HPLC (Figure 3C-3D, Figure S3). Some clones with both sub-region 4 alleles deleted exhibited increased proportion of HbF-expressing cells (up to 40% for cells derived from HUDEP-2, but not for lines from H2.1, Figure 3C). However, other deletion clones did not exhibit any apparent increase in HbF. A similar but reduced effect was observed for Region 9. Overall, we observed extreme variation in HbF-expressing cells between clonal cell lines with the same genotype, whether derived from polyclonal HUDEP-2 line or the isogenic H2.1 sub-clone.

We attempted to validate our flow cytometry results by HPLC (Figure 3D, traces in Figure S3). Two sub-region 4 homozygous knockout clones derived from HUDEP-2 exhibited high proportions of HbF-expressing cells. However, all other clones, including all region 4 knockout clones derived from the H2.1, did not exhibit high HbF. These results suggest that sub-regions 4 and 9 are not strictly necessary for γ-globin silencing in HUDEP-2 cells, and highlight significant clonal differences in proportions of HbF-expressing HUDEP-2 cells.

Finally, we assessed whether selected PRR or PRR sub-region deletion genotypes can alter γ-globin expression in erythroblasts derived from human CD34+ hematopoietic stem/progenitor cells (HSPCs). To obtain data from clonal erythroid cells we used a combination of Cas9 RNP editing of CD34+ HSPCs and an erythroid colony assay. Each BFU-E arising from an edited HSPC can readily be genotyped using PCR and amplicon NGS, and globin gene expression can be assessed using RNA-seq (Figure 4A) *(17)*.

**Figure 4:**
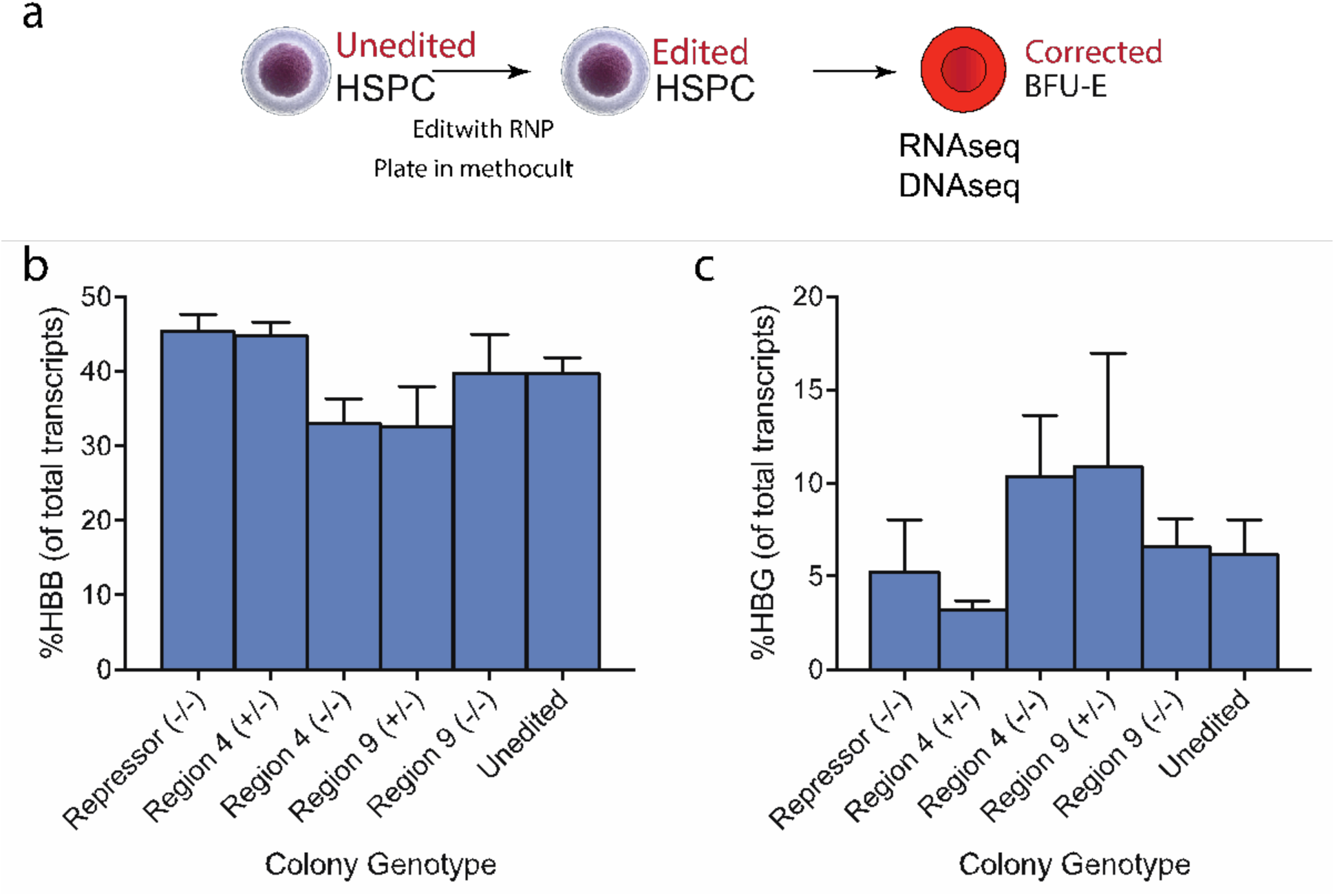
Genotype-phenotype relationships in clonal BFU-E colonies derived from edited hematopoietic stem/progenitor cells. A) Workflow depicting erythroid colony formation assay using edited HSPC. HSPC were edited using RNPs targeting the desired region, plated in methocult, and colonies were grown for 14 days. Erythroid colonies were picked and genotyped by next-generation sequencing; selected colonies were assayed for globin expression by RNA-seq. B) HBB expression in erythroid colonies after gene editing, determined by RNA-seq. C) γ-globin expression in erythroid colonies after gene editing, determined by RNA-seq.

As in the experiments described above, we used pairs of Cas9 RNPs to target deletion of the 3 regions of greatest interest: the full 1.7 kb PRR, and sub-regions 4 and 9. HSPCs were editing using our published protocols *(26)*, and plated in MethoCult Express (StemCell Technologies) to form colonies (Figure 4A). The resulting erythroid colonies (BFU-E) were individually removed for genotyping and RNA-Seq. In general globin mRNA made up ∼60% of all transcripts, consistent with late-stage erythroid maturation. ß-globin (HBB) expression was largely unaltered across the 5 genotypes (PRR (-/-), Sub-region 4 (-/-) and (+/-), Sub region 9 (-/-) and (+/-)) (Figure 4B). γ−globin (HBG1+HBG2) expression was highly variable between clonal colonies, ranging from 0.5% to 26% of all transcripts, with ß-globin (HBB) expression ranging from 23% to 50% of all transcripts (Figure 4C). There was no clear association between PRR deletion genotype and globin expression phenotype, indicating that the neither the full PRR nor either of the selected sub-regions is required for γ-globin silencing in primary human erythroblasts.

## Discussion

Here we explored a potential approach to activate expression of γ-globin in adult erythroid cells via targeted ablation of a putative repressor region (PRR) upstream of *HBD*, identified from analysis of naturally occurring HPFH deletions (Figure S1) *(19)*. We used the human erythroblast cell line HUDEP-2, along with primary human CD34+ HSPCs, to ask if this region is necessary for maintenance of γ-globin silencing. Although our initial screens in bulk populations of cells suggested a role for the PRR in γ-globin silencing, we found that deletion of the entire region selected sub-regions were not consistently associated with γ-globin expression in clonal HUDEP-2 cell lines or in erythroid colonies derived from edited CD34+ HSPCs. We conclude that deletion of the PRR is not sufficient for stable up-regulation of γ-globin, and thus the PRR is not a suitable target for a gene editing therapy.

While initial experiments with Cas9 RNPs that drive deletion of the PRR or its sub-regions found that some RNPs were associated with increased numbers of HbF-expressing cells in the polyclonal HUDEP-2 cell line, a more detailed examination with clonal sublines carrying defined deletions did not support the preliminary result. This discrepancy, and the broad variation in HbF expression among clonal sublines of HUDEP-2, suggest that HUDEP-2 cells exhibit spontaneous but unstable expression of γ-globin. It is not clear why electroporation of some Cas9 RNPs induced HbF expression in subpopulations of HUDEP-2 cells, but the analysis of clones militates against this effect being a valid indication that the PRR is a γ-globin silencer. The result is not without value, since it indicates that caution should be used in interpreting results from clonal HUDEP-2 sublines. Deriving edited clones at required scale can be challenging. Instead, model cell lines such as HUDEP-2 may be dispensed with entirely: efficient NHEJ-mediated gene editing of HSPCs is feasible with Cas9 RNP electroporation *(26)*, and the relatively few cells in clonal BFU-E colonies can readily be genotyped by NGS and phenotyped with RNA-seq *(16, 17)*.

The PRR we have studied was identified as a putative γ-globin silencer on the basis of two lines of evidence *(19)*. One was the comparison of multiple large deletions in the ß-globin locus: large deletions that do not include the PRR cause ß-thalassemia, while other deletions that do include this region confer an HPFH phenotype. The second was a ChIP-Seq analysis that identified binding of BCL11A within the PRR *(19)*. BCL11A is well documented as a regulator of developmental γ-globin silencing *(28)*, and so this result seemed consistent with the idea that the region was required for developmental silencing. However a more recent study that used a different method to map BCL11A binding sites did not identify binding within the PRR *(29)*.

As this manuscript was in preparation a new study exploring deletions of genomic regions encompassing the PRR was reported using similar methods *(18)*. Interestingly, researchers found that deletion of the 7.2 kb Corfu deletion, or a shorter 3.5 kb region, both including the 1.72 kb PRR, did not lead to significant fetal hemoglobin re-expression in HUDEP-2 cells. Instead a larger 13.2 kb deletion that encompassed the adult ß-globin promotor and led to 3-fold drop in relative ß-globin expression did lead to robust γ-globin expression, similar to HPFH-1 deletion using CRISPR/Cas9 in HSPCs *(17)*.

The possibility remains that regulatory complexes bound to the PRR cooperate with other regions within or downstream of HBB to mediate γ-globin silencing. Previous studies have shown that deletions including both the PRR and all or part of ß-globin gene, along with other downstream elements, can induce γ-globin *(17)*. However the safe and efficient induction of these more extended deletions would be challenging in a clinical setting.

## Acknowledgements

This work used the Vincent J. Coates Genomics Sequencing Laboratory at UC Berkeley, supported by NIH S10 OD018174 Instrumentation Grant. Synthetic RNAs for studies presented in Figure 3D were provided by Synthego, Inc. This work was supported by the Li Ka Shing Foundation (J.E.C.), the Heritage Medical Research Institute (J.E.C.), the California Institute of Regenerative Medicine (M.A.D., J.E.C.), the Siebel foundation (M.A.D.), the Lam foundation (J.Chung), and the Jordan Family Foundation (D.K.M.).

## Materials and Methods

*HUDEP-2 Cells and cell culture.* HUDEP-2 cells were cultured and differentiated according to published protocols *(23)*, with the exception that doxycycline was included in *both* differentiation and expansion media. Expansion medium: 1 μM dexamethasone, 1 μg/mL doxycycline, 50 ng/mL stem cell factor (Peprotech, Inc.), 5 U/mL EPO (Amgen, pharmaceutical grade), in SFEM medium (Stem Cell Technologies). Differentiation medium: 5% human serum (Sigma Aldrich), 2 IU/mL heparin (Sigma Aldrich), 10 μg/mL Insulin (Sigma Aldrich), 5 U/mL EPO (Amgen), 500 μg/mL holo-transferrin (Sigma Aldrich), 1 μM mifepristone (Sigma Aldrich), 1 μg/mL doxycycline in IMDM media (with Glutamax, Gibco, Inc.). Differentiation medium was prepared with one week of use and sterile-filtered using a 0.22 μm filter. To differentiate HUDEP-2 and H2.1 cells from expansion culture, cells were pelleted (300 × g, 5 minutes), thoroughly decanted, re-suspended in differentiation medium at a density of <1,000,000 cells/mL, and cultured for 5 days before analysis for hemoglobin expression by flow cytometry or HPLC.

*Synthesis of Cas9 RNPs*. Cas9 RNP component synthesis and assembly was carried out based on published work, and are available online at https://www.protocols.io/groups/igi/protocols *(26, 30)*. Cas9 was prepared by the UC Berkeley Macro Lab using a published protocol *(30)*. Cas9 was stored and diluted in sterile-filtered Cas9 Buffer (20 mM HEPES pH 7.5, 150 mM KCl, 1 mM MgCl_2_, 10% glycerol, 1 mM TCEP). sgRNA was synthesized by assembly PCR and *in vitro*-transcription. A T7 RNA polymerase substrate template was assembled by PCR from a variable 59 nt primer containing T7 promotor, variable sgRNA guide sequence, and the first 15 nt of the non-variable region of the sgRNA (T7FwdVar primers, 10 nM, Table S3), and an 83 nt primer containing the reverse complement of the invariant region of the sgRNA (T7RevLong, 10 nM), along with amplification primers (T7FwdAmp, T7RevAmp, 200 nM each). Phusion high-fidelity DNA polymerase was used for assembly (New England Biolabs, Inc.). Assembled template was used without purification as a substrate for *in vitro* transcription by T7 RNA polymerase using the HiScribe T7 High Yield RNA Synthesis kit (New England Biolabs, Inc.). Resulting transcriptions reactions were treated with DNAse I, and purified either by with a 5X volume of homemade SPRI beads (comparable to Beckman-Coulter AMPure beads), and eluted in DEPC-treated water. sgRNA concentration was determined by fluorescence using the Qubit RNA BR assay kit (Life Technologies, Inc). Cas9 RNP was assembled immediately prior to electroporation of target cells (see below). To electroporate a 20μL cell suspension (see below) with Cas9 RNP, a 3.75 μL solution containing a 1.2-1.3X molar excess of sgRNA in Cas9 buffer was prepared. A 3.75 μL solution containing 75 pmol purified Cas9 in Cas9 buffer was prepared and added to the sgRNA solution slowly over ∼30 seconds, and incubated at room temperature for >5 minutes prior to mixing with target cells. For electroporation with pairs of Cas9 RNPs, half the amounts and volumes above were used to assemble RNPs, which were then mixed after assembly. For studies depicted in Figure 2E (saturating mutagenesis of PRR sub-region 4), RNA was generously provided by Synthego, Inc. For electroporation of HSPCs (Figure 4), synthetic sgRNA was purchased from Synthego, Inc. Synthesis of RNPs from synthetic sgRNA was accomplished in an analogous manner, using stocks of dried RNA hydrated to 45 μM in water or 0.5X TE. *Electroporation of HUDEP-2 Cells with Cas9 RNP using the Lonza 4d electroporator.* HUDEP-2 cells were cultured in expansion medium prior to editing to mid-log phase (200,000-1 million cells per mL of culture). To electroporate, 100,000-200,000 cells were spun at 300xg for 5 minutes, and resuspended in 20 μL of Lonza P3 solution and 7.5 μL of Cas9 RNP solution prepared as described above, before transfer of the entire 27.5 μL mixture to a small-scale Lonza S electroporation cuvette. Cells were electroporated using Lonza 4d code DD100. Cells were cultured a minimum of 2 days before differentiation and phenotyping analysis. *Generation of clonal HUDEP-2 sublines.* Gene-edited HUDEP-2 and H2.1 cells were cloned by limiting dilution into 96 well plates. After ∼2 weeks the cultures were passaged and half of the cells were removed for DNA extraction and PCR (see above). Genotyping was initially performed with standard PCR and agarose gel electrophoresis using primers for the indicated deletion (table S3). Clones with desired genotypes by PCR were confirmed with next-generation sequencing (see below).

*Assay of HbF-positive HUDEP-2 cells by intracellular flow cytometry.* FACS for HbF was adapted from existing protocols *(24)*. Differentiated or undifferentiated HUDEP-2 cells were pelleted by centrifugation at 500 × g for 5 minutes, washed with PBS/0.1% BSA, pelleted again and resuspended in 200 μL 0.05% glutaraldehyde (freshly prepared from evacuated ampules of 20% glutaraldehyde from Sigma Aldrich) in PBS and incubated for 10 minutes to fix the cells. Cells were pelleted at 600 × g for 5 minutes and resuspended in 200 μL of 0.1% Triton X-100 in PBS/0.1% BSA and incubated for 10 minutes to permeabilize cells, and pelleted at 600 × g once again before resuspension in 50 μL of anti-fetal hemoglobin-FITC antibody at a 1-10 dilution in PBS/0.1% BSA (BD Biosciences, Inc. clone 2D12) and incubated for 20 minutes to stain cells. After staining, cells were washed three times in PBS/0.1% BSA before analysis by flow cytometry.

*Quantification of relative hemoglobin expression with HPLC.* HPLC analysis for hemoglobin expression in differentiated cells was run essentially as described in previous work *(31)*. Hemolysates for HPLC were prepared from 1-5 million differentiated cells pelleted and resuspended at 100,000 cells/μL in hemolysate reagent (Helena Laboratories), incubated for 10 minutes at room temperature, and clarified by centrifugation. 10 μL of lysate was applied to the HPLC column for hemoglobin analysis. For HPLC a PolyCAT A column 3.54 (PolyLC, Inc.) was used with mobile phase A for loading (20 mM Bis-tris, 2 mM NaCN, pH6.8) and B for elution (20 mM Bis-tris, 2 mM NaCN, 200 mM NaCl, pH 6.9), flow-rate 1.5 mL/min, and detection of hemoglobin by absorbance at 415 nm, and gradient from 0% to 100% mobile phase B over 30 minutes to separate hemoglobins. Hemoglobins were identified based on mobility using a separate FASC hemoglobin standard (Trinity biotech).

*Electroporation of CD34+ HSPCs with Cas9 RNPs.* HSPC electroporation was as described *(26)*. CD34+ HSPCs (G-CSF mobilized, Allcells, Inc.) were thawed and cultured in SFEM (StemCell Technologies, Inc.) with CC110 cytokines (Stem Cell Technologies) for 2 days at a density of <500,000 cells/mL before editing. 100,000-200,000 HSPCs were re-suspended in 20 μL of P3 solution and 7.5 μL of Cas9 RNP or RNP pairs assembled as described above. Because ssDNA reduces viability when gene editing HSPCs, no “stapler” ssDNAs were used to edit HSPCs. For HSPCs, only synthetic sgRNA was used (Synthego, Inc.). HSPCs in P3 solution with RNP were electroporated using code ER100 on a Lonza 4d electroporator. After electroporation, cells were layered with 75 μL of SFEM with CC110 and incubated for 5 minutes at room temperature before transfer to culture in SFEM with CC110. Cells were cultured for 1 day before plating on MethoCult Express at 250 cells/well in a 6 well plate. A subset of the culture (∼50,000 cells) was taken to confirm deletions by PCR. After 14 days, individual BFU-E were picked and dispersed in 100 μL of PBS. Half of the suspension was taken for genotyping by genomic PCR and next-generation sequencing, and the other half was stored at -80. Colonies found to have the desired genotypes by genomic DNA PCR were thawed and prepared for RNA-seq, and had their genotypes confirmed by next-generation sequencing before completion of RNA-seq data analysis (see below).

*RNA-seq of erythroid colonies* Total RNA was isolated from individual BFU-E with the ArrayPure Nano-scale RNA purification kit (Epicentre), and converted into cDNA following the Smart-seq2 protocol *(32)*. The cDNA was fragmented (median size of 170 bp) and ligated into Illumina sequencing libraries with the KAPA HyperPlus kit (KAPA Biosystems). Libraries were sequenced on an Illumina HiSeq4000 apparatus to produce single-ended reads of 50 nucleotides.Reads were aligned with kallisto version 0.43.1 *(33)* against a reference sequence consisting of the main isoforms of each globin gene at the alpha-and beta-globin loci. The following Ensembl transcript IDs were used: ENST00000320868.9 (HBA1), ENST00000251595.10 (HBA2), ENST00000199708.2 (HBQ1), ENST00000356815.3 (HBM), ENST00000354915.3 (HBZP1), ENST00000252951.2 (HBZ), ENST00000335295.4 (HBB), ENST00000380299.3 (HDB), ENST00000454892.1 (HBBP1), ENST00000330597.3 (HBG1), ENST00000336906.4 (HBG2), ENST00000292896.2 (HBE1).

Globin transcripts have high levels of sequence identity; to evaluate the ability of kallisto to correctly assign a read to the transcript from which it was derived, we carried out simulations using computationally generated 50-base reads. The globin transcripts listed above were mixed in known proportions that simulate relative transcript levels in erythrocytes (HBA-1: 0.7, HBA-2: 0.7, HBM: 0.01, HBB: 1.00, HBD: 0.04). HBG-1 and HBG-2 were varied between 0.005 and 0.10 to simulate the effect of varying levels of gamma-globin expression in a mixture. Sequence segments of 50 nucleotides in length were sampled randomly from this mixture and from its reverse complement (number of samples = 1,000,000), and aligned as described above. The random sampling procedure was repeated 10 times for each mixture. For each alignment result, we calculated the mean ratio of observed over expected counts for each transcript, which we used as adjustment factors for the counts reported by kallisto after aligning real data; the difference in transcript counts before and after adjustment ranged from -1% to 0.5%. The computed adjustment factors are listed below:

HBA-1 1.0397665108
HBA-2 0.8890515518
HBM 0.8195218055
HBB 1.0401825121
HBD 1.1146242591
HBG-1 1.321849175
HBG-2 1.0079414563

*T7 Endonuclease assay.* To select optimal gRNAs, genomic cutting using Cas9 RNPs was estimated using T7 endonuclease I assay. Genomic DNA was extracted from edited cells (HUDEP-2 or CD34+ HSPC) using QuickExtract solution (Epicentre, Inc.) to a density of >2,500 cells/μL. A PCR amplicon of the targeted region was generated using GXL Primestar Polymerase (Takara, Inc.), 30 PCR cycles, and manufacturer’s instructions, at a final volume of 50 μL using 5 μL of extracted genomic DNA as template. 7 μL of unpurified PCR was mixed with water and 1X NEB buffer 2 to a final volume of 10 μL before re-annealing in a thermal cycler (95 degrees for 5 minutes, cool to room temperature over 10 minutes). 1 μL of T7 endonuclease I was added (NEB, Inc.) and reaction was digested for 30 minutes at 37°C before visualization on a 2% agarose gel or 4-20% polyacrylamide TBE gel (Life Technologies, Inc.). Cutting was compared to PCR amplicons from untreated cells.

*Genotyping of HSPC colonies and HUDEP-2 clonal cell lines edited at the PRR and sub-regions by PCR amplicon deep sequencing (NGS).* Genomic DNA preparation and PCR were performed identically as for T7 endonuclease digest above, except that the PRR primers were used for all samples (see supplemental text for unedited amplicon sequence). The 2.3 kb amplicon was purified, fragmented to an average length of ∼350 bp using a Covaris instrument, and prepared for NGS using the Illumina TruSeq Nano HT kit. Libraries (1 for each colony or clonal cell line) were sequenced on an Illumina MiSeq (2×300 paired-end sequencing). Demultiplexed and adaptor-trimmed reads were aligned to the reference amplicon sequence, and the genotype determined by visualization in IGV viewer software.

**Figure S1.**
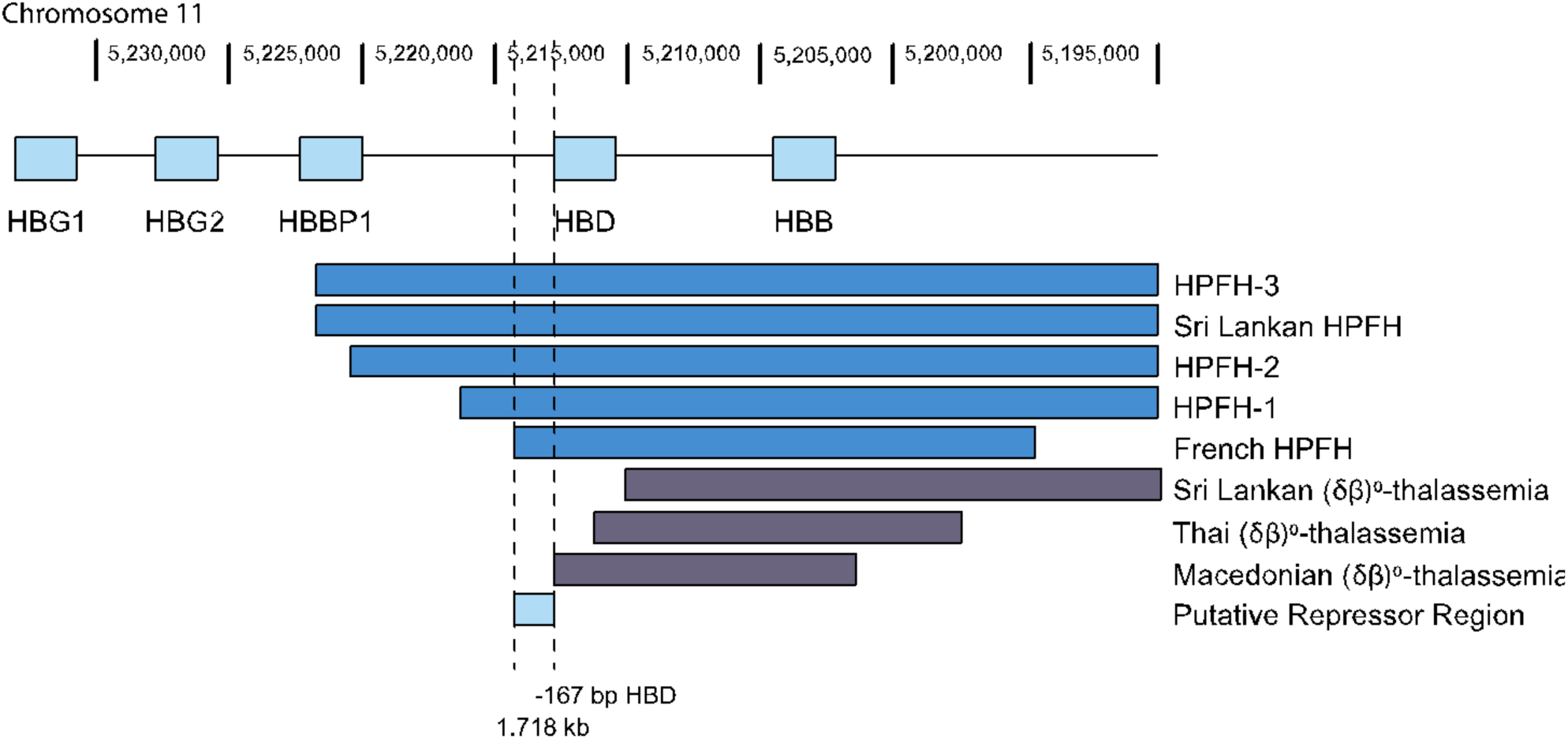
Defining a putative repressor region from naturally-occurring deletion HPFH mutations, based upon *(13)*. Mapping of observed mutations associated with beta thalassemia (grey bars) and observed mutations associated with HPFH (blue bars), it appears that deletion of a 1.72 kb region starting 167 bp upstream of the HBD gene is associated with an HPFH phenotype, while deletions that retain this region are associated with a ß-thalassemia phenotype.

**Figure S2.**
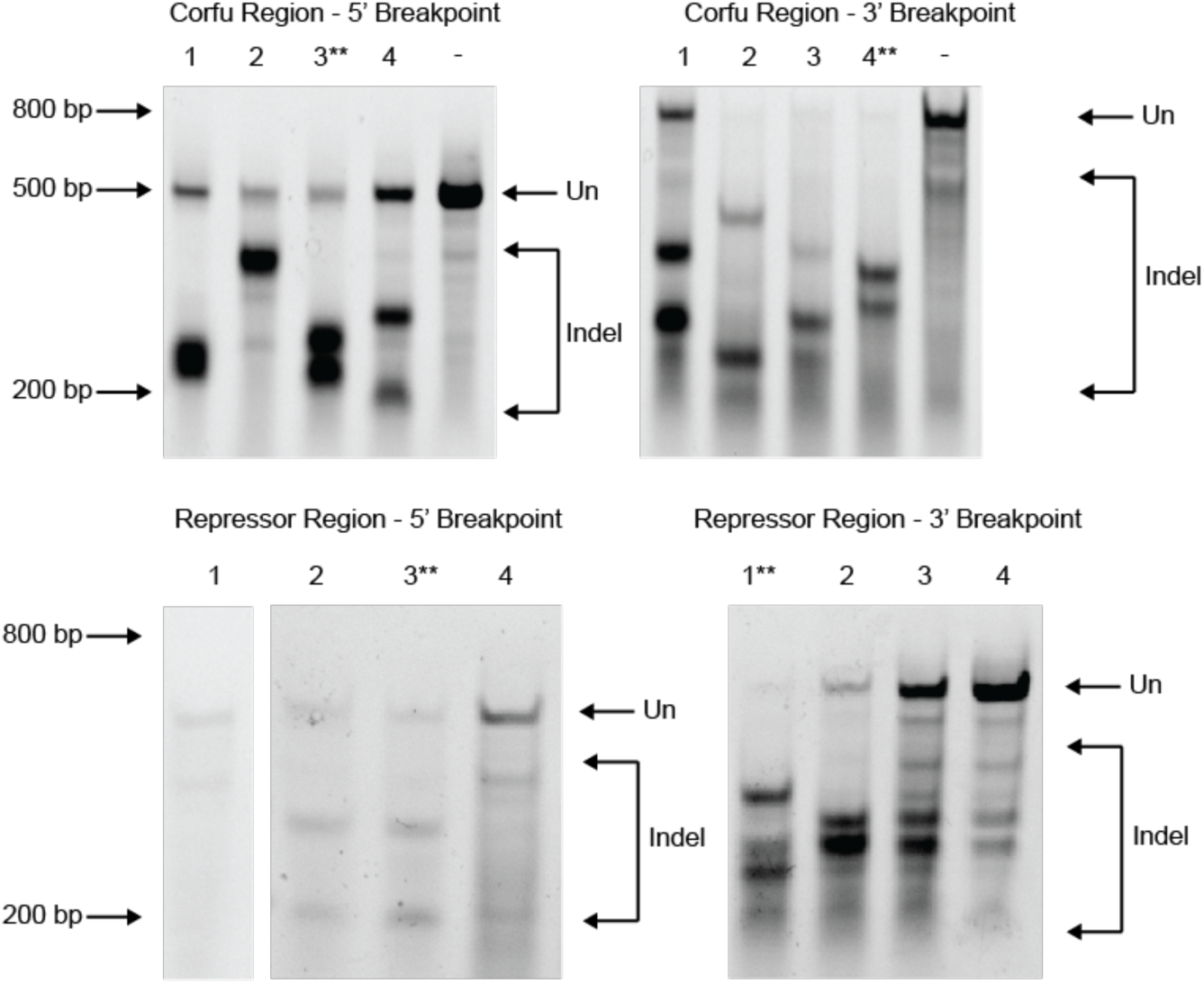
Development of RNP pairs deleting two HPFH-associated regions. Guides targeting the 4.5 kb Corfu deletion were electroporated in RNPs and assessed by T7 endonuclease digestion of PCR amplicons. Guides with the most efficient targeting as measured by T7 digestion were used as pairs to target the full regions. For the Corfu region, guide 3 on the 5’ side and guide 4 were selected (top row). For the PRR, guide 3 on the 5’ end (1.8 kb upstream of HBD) and guide 1 on the 3’ (adjacent to HBD) were selected (middle row).

**Figure S3.**
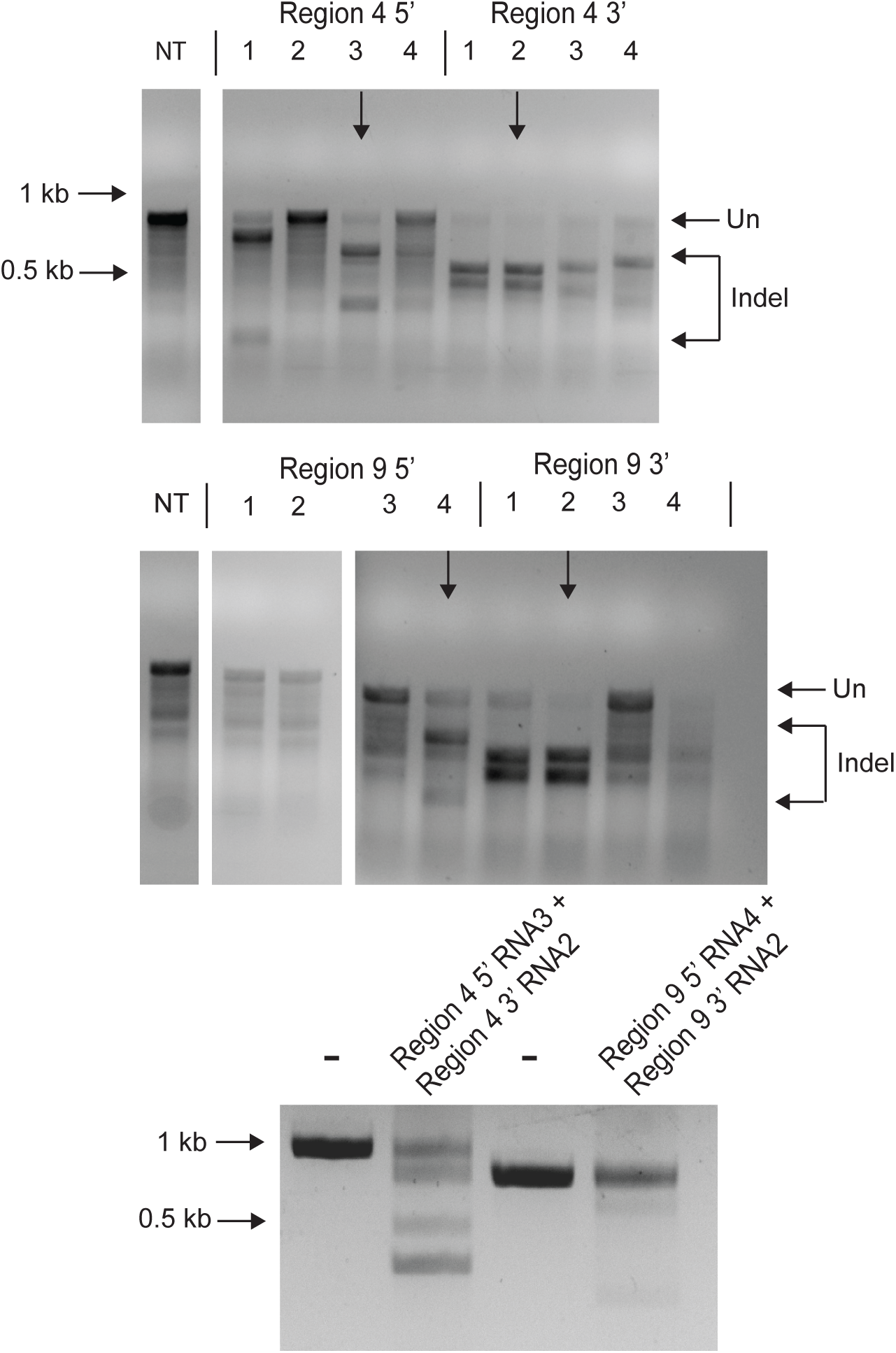
Development of RNP pairs deleting sub-regions within the putative repressor region (Related to figure 3). 4 guides targeting each breakpoint were synthesized by *in vitro* transcription, formed into RNPs and delivered to subcloned HUDEP-2 cells by electroporation. 48 hours after electroporation, cells were harvested and the targeted regions were amplified by genomic PCR and edited in individuals RNPs was assessed by T7 endonuclease I digest (above). Optimized paired RNPs were co-delivered by electroporation and deletion was observed with genomic PCR (below).

**Figure S4.**
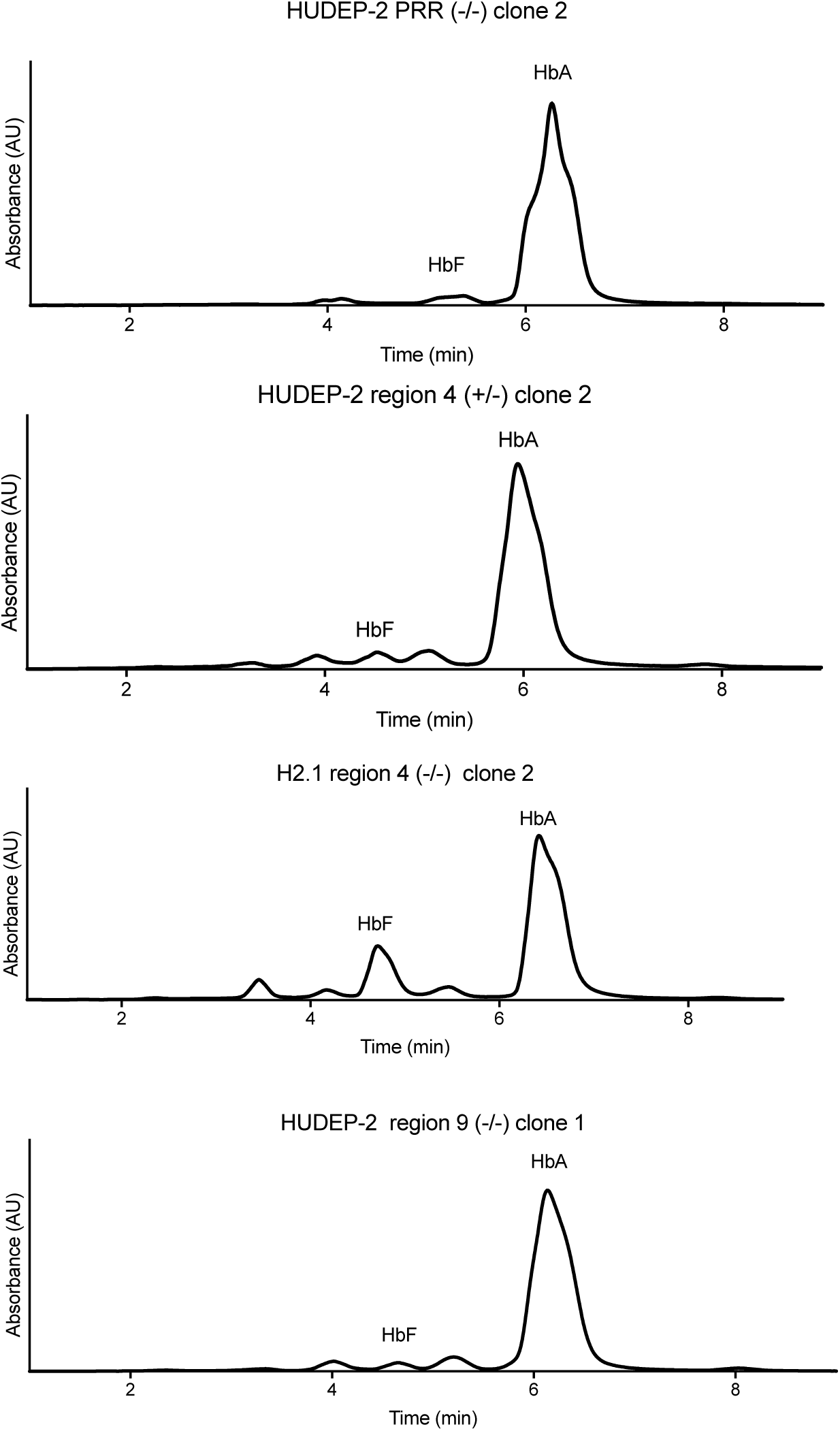
Representative of HPLC traces of HUDEP-2 clones with the full PRR or either of sub-regions 4 or 9 deleted, as indicated. Hemoglobins were identified with FASC standards run separately.

**Table S1.**
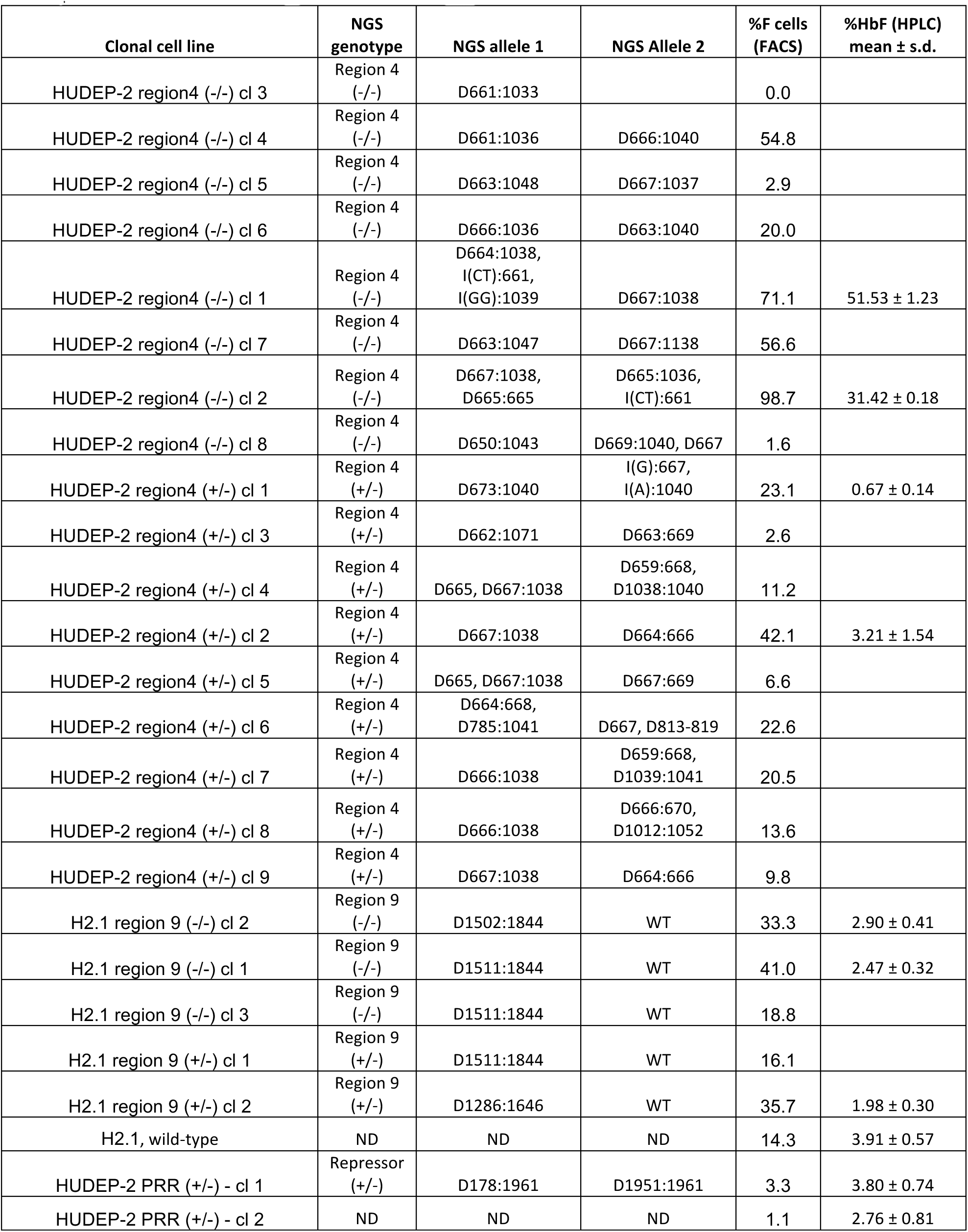
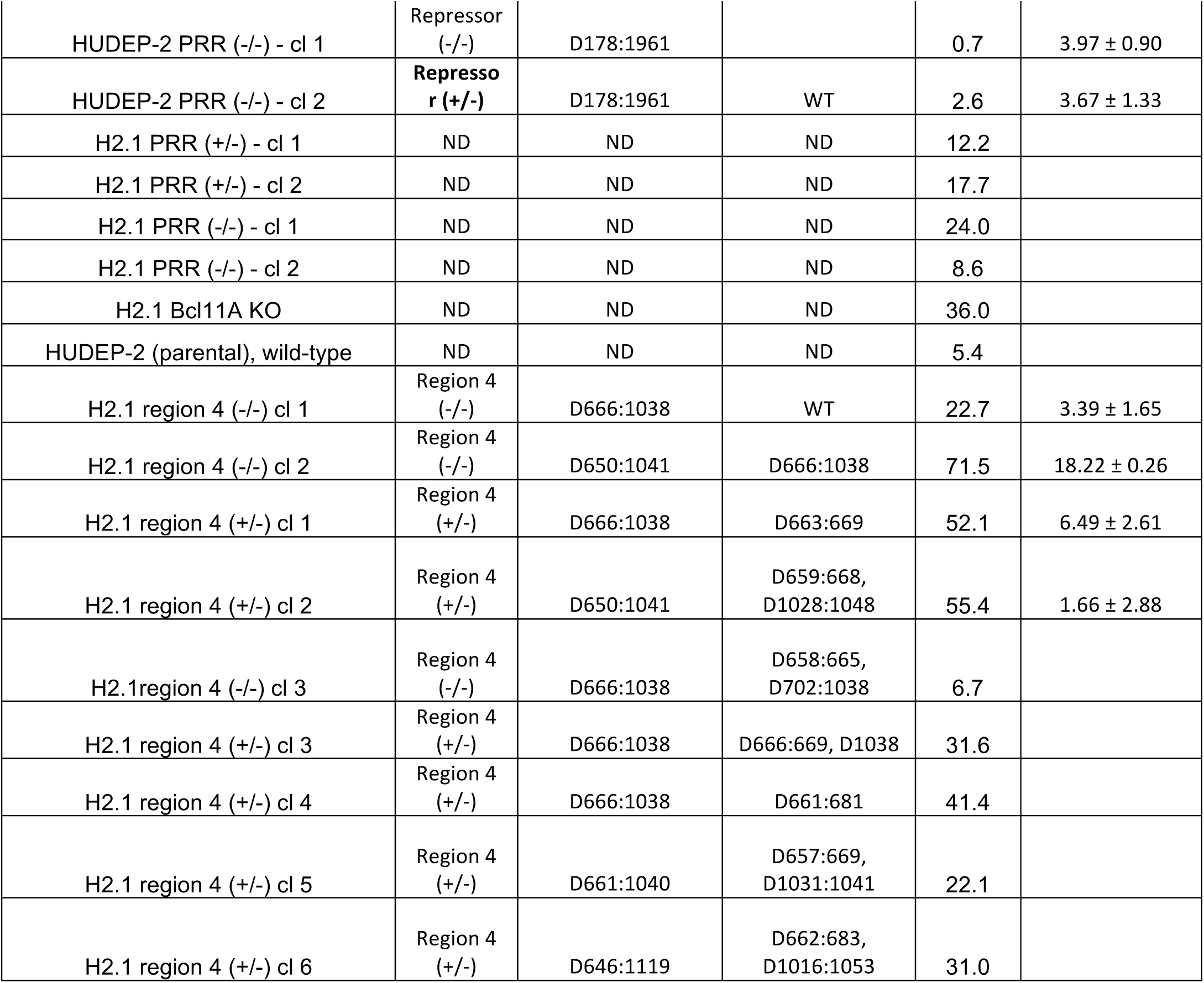
NGS genotyping results of clonal cell lines derived in this study. NGS genotypes are Insertions (“I”) or deletions (“D”) of the indicated nucleotides at the indicated positions within the repressor region NGS amplicon (see below).

**Table S2.** NGS genotyping results of clonal CFU-E colonies derived from HSPCs after editing with Cas9 RNP pairs programming deletion of the three regions and sub-regions of interest in this study. NGS genotypes are Insertions (“I”) or deletions (“D”) of the indicated nucleotides at the indicated positions within the repressor region NGS amplicon (see supplemtal text below). Globin transcript abundance by RNA-seq (as %of all transcripts) is indicated.

| NGS genotype | NGS allele 1 | NGS Allele 2 | %HBG1<br>+HBG2 | %HBB | %HBD | %HBA |
| --- | --- | --- | --- | --- | --- | --- |
| PRR (-/-) | D178:1961 |  | 7.88% | 45.20% | 1.67% | 44.51% |
| PRR (+/-) | D184:1979,189(TC->AA) | D1962 (T) |  |  |  |  |
| PRR (-/-) | D172:1962 | D181:1954,<br>1954ATTCCAC-<br>>CTCCAGATC | 1.91% | 48.04% | 1.21% | 48.08% |
| PRR (+/-) | D178:1963 | D1963, D180-183 |  |  |  |  |
| PRR (+/-) | D180:1962 | D1962 (T) |  |  |  |  |
| PRR (-/-) | D178:1961 |  | 14.79% | 38.42% | 1.31% | 44.33% |
| PRR (-/-) | D178:1961 | D178:1961<br>I(1961):TGT | 0.54% | 49.80% | 1.44% | 47.75% |
| PRR (-/-) | D178:1961 |  | 1.53% | 46.73% | 1.88% | 49.23% |
| Region 4 (+/-) | D661:1034,<br>1034CACCA:ctctg | I665:G | 2.56% | 45.88% | 2.27% | 48.45% |
| Region 4 (+/-) | D662:1089 | D663:669 | 3.34% | 47.24% | 2.39% | 46.11% |
| Region 4 (+/-) | D661:1034,<br>1034CACCA:ctctg | D664-667 | 3.95% | 41.81% | 1.45% | 52.27% |
| Region 4 (+/-) | D666:1039 | D1024:1061 |  |  |  |  |
| Region 4 (+/-) | D666:1038 | D666:674 |  |  |  |  |
| Region 4 (+/-) | D667:1038 | Unedited |  |  |  |  |
| Region 4 (-/-) | D666:682,691:1543 | D656:986 | 7.33% | 36.29% | 0.88% | 54.91% |
| Region 4 (-/-) | D661:1034 |  | 13.60% | 30.42% | 1.02% | 54.51% |
| Region 9 (+/-) | D1517:1850 | D1844 |  |  |  |  |
| Region 9 (+/-) | D1512:1844 | I1841:TC |  |  |  |  |
| Region 9 (+/-) | D1517: S1512:5:TAGA | Unedited |  |  |  |  |
| Region 9 (-/-) | D1517:1849, Mut1512-6:TAGAG |  | 15.03% | 23.38% | 1.45% | 58.75% |
| Region 9 (-/-) | D1512:1844 |  | 2.70% | 42.15% | 2.23% | 52.05% |
| Region 9 (-/-) | D1508:1843 ,I1507:GC,<br>M1844:A | D1508:1843 ,S1844:A | 26.04% | 24.20% | 1.09% | 47.75% |
| Region 9 (-/-) | D1512:1844 |  | 0.20% | 41.31% | 1.71% | 56.26% |
| Region 9 (+/-) | D1512:1844 | D1845:1846 | 4.08% | 46.26% | 2.09% | 46.96% |
| Region 9 (+/-) | D1511:1845 | D1845:1849 | 7.34% | 43.54% | 2.67% | 45.57% |
| Region 9 (+/-) | D1512:1844 | M1530:T-G,D1841:2 | 8.72% | 30.19% | 0.45% | 60.38% |
| wild-type | Unedited |  | 11.97% | 33.45% | 1.28% | 52.34% |
| wild-type | Unedited |  | 4.78% | 39.05% | 1.72% | 53.71% |
| wild-type | Unedited |  | 5.27% | 42.32% | 2.46% | 49.07% |
| wild-type | Unedited |  | 7.94% | 41.44% | 1.83% | 47.99% |
| wild-type | Unedited |  | 26.46% | 24.10% | 0.99% | 47.49% |
| wild-type | Unedited |  | 1.50% | 43.56% | 1.45% | 52.96% |
| wild-type | Unedited |  | 13.38% | 37.09% | 1.92% | 46.68% |

**Table S3.**
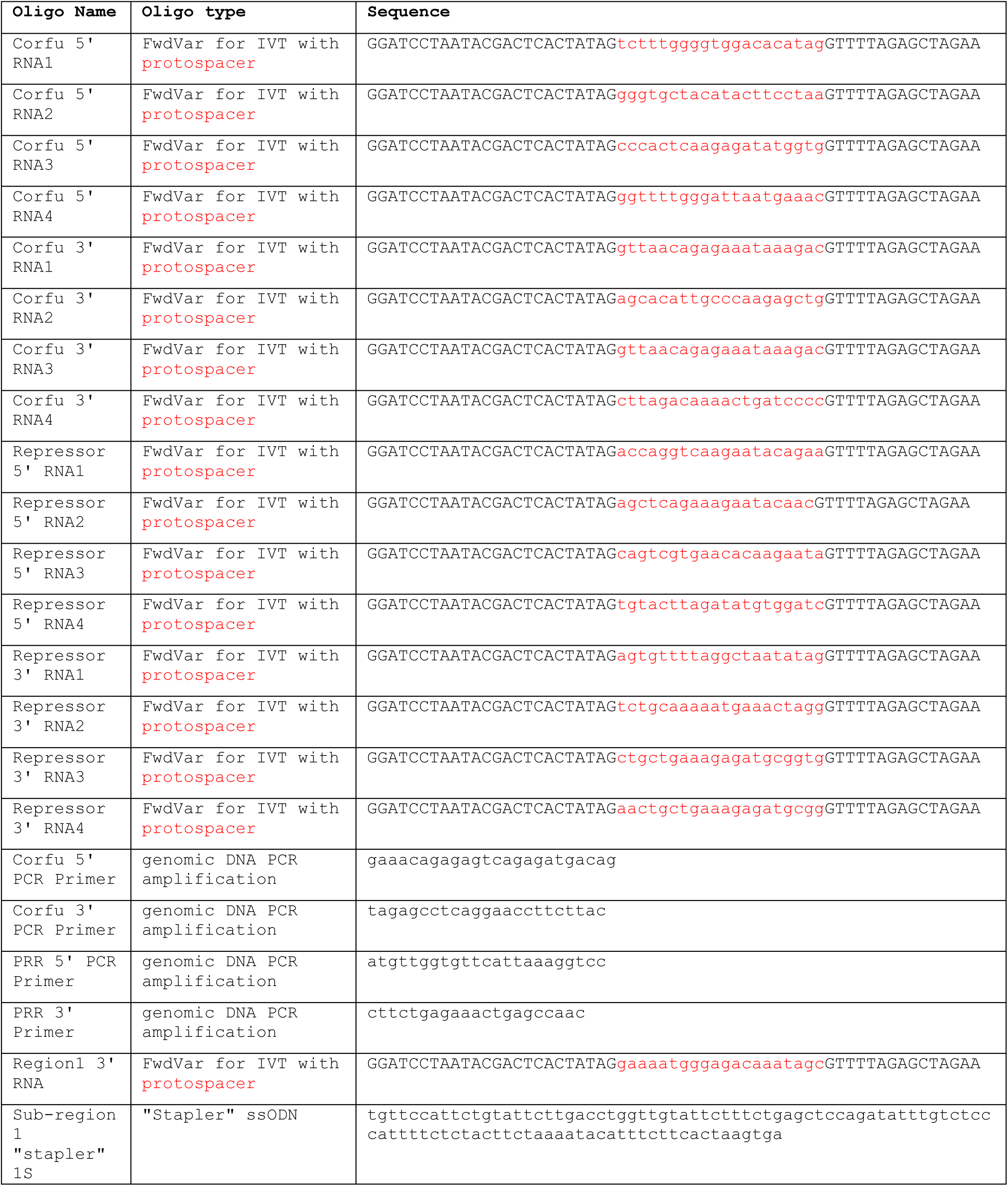
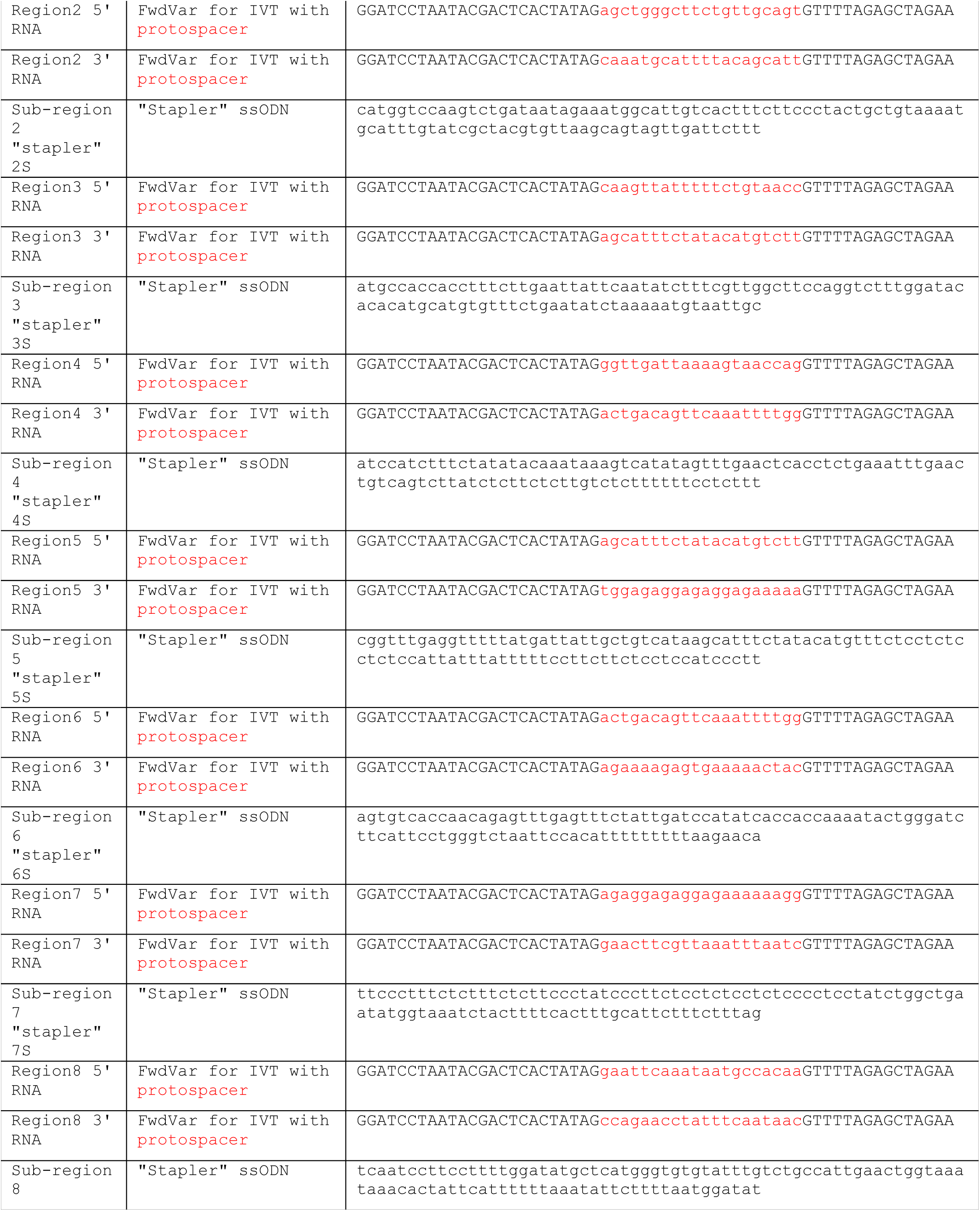
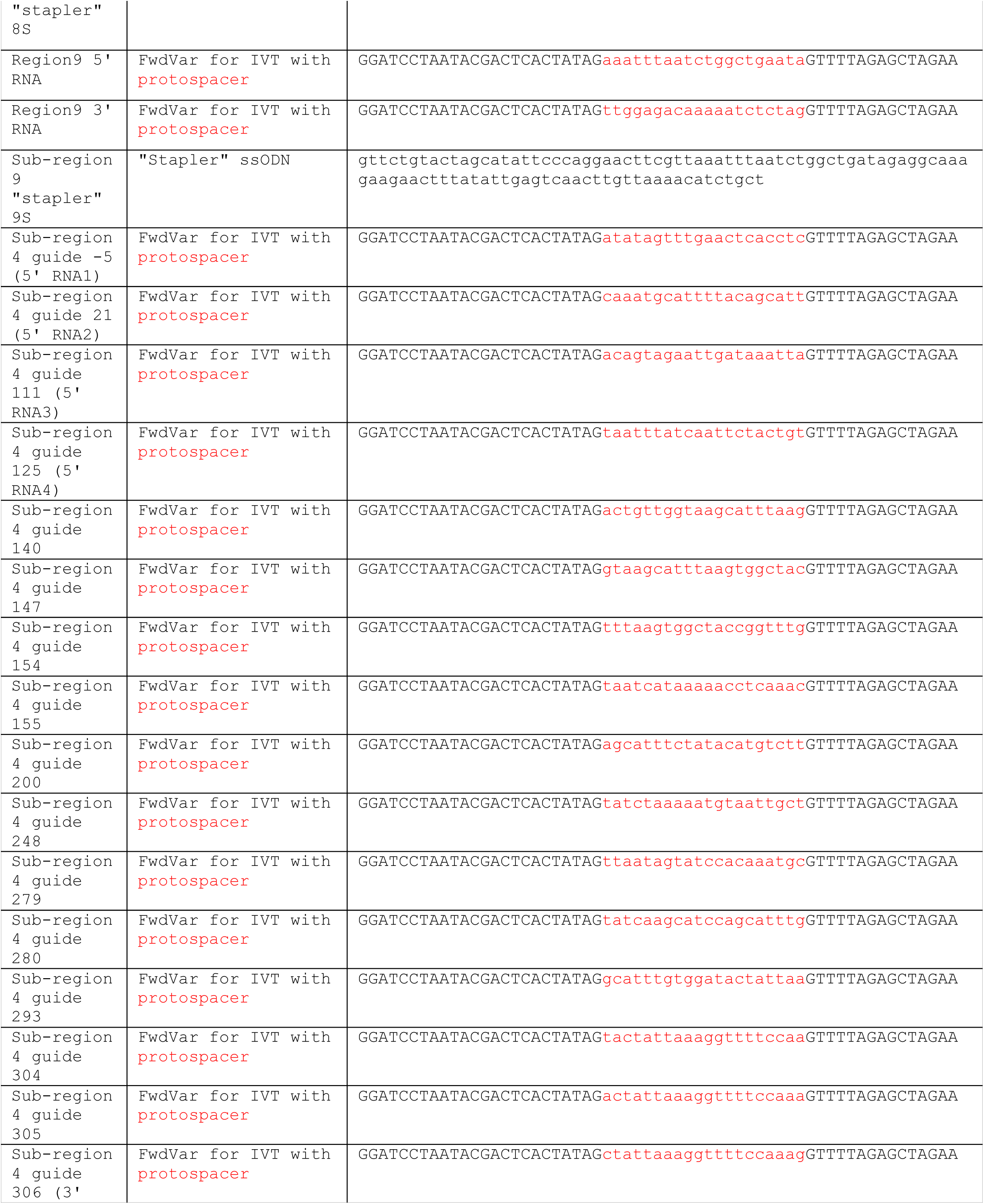
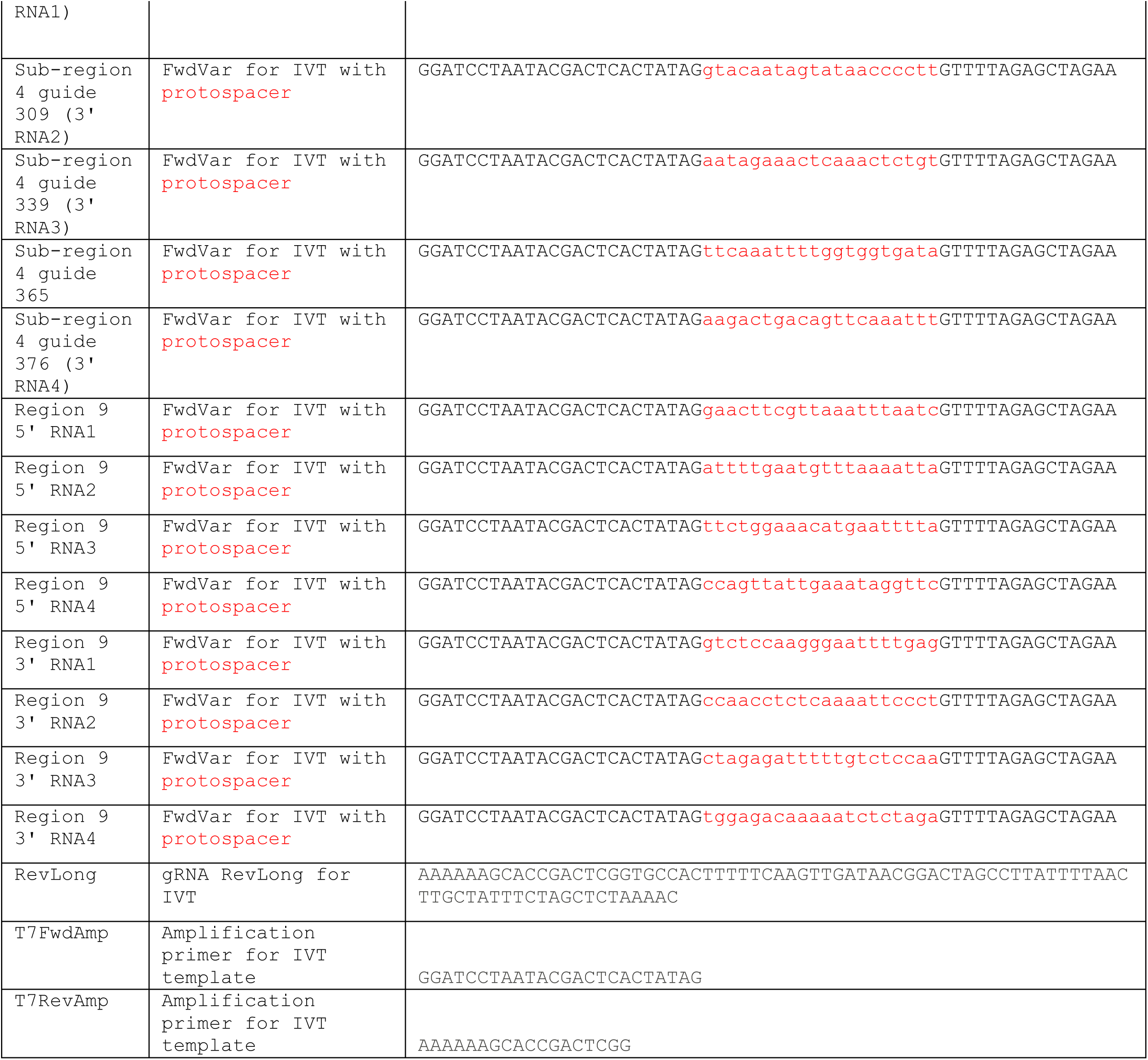
DNA Oligonucleotides used in this study, including gRNA protospacer sequences where indicated.

Supplemental Information

I. Repressor region PCR amplicon, used as a reference for NGS analysis of clonal cell lines and HSPC colonies.

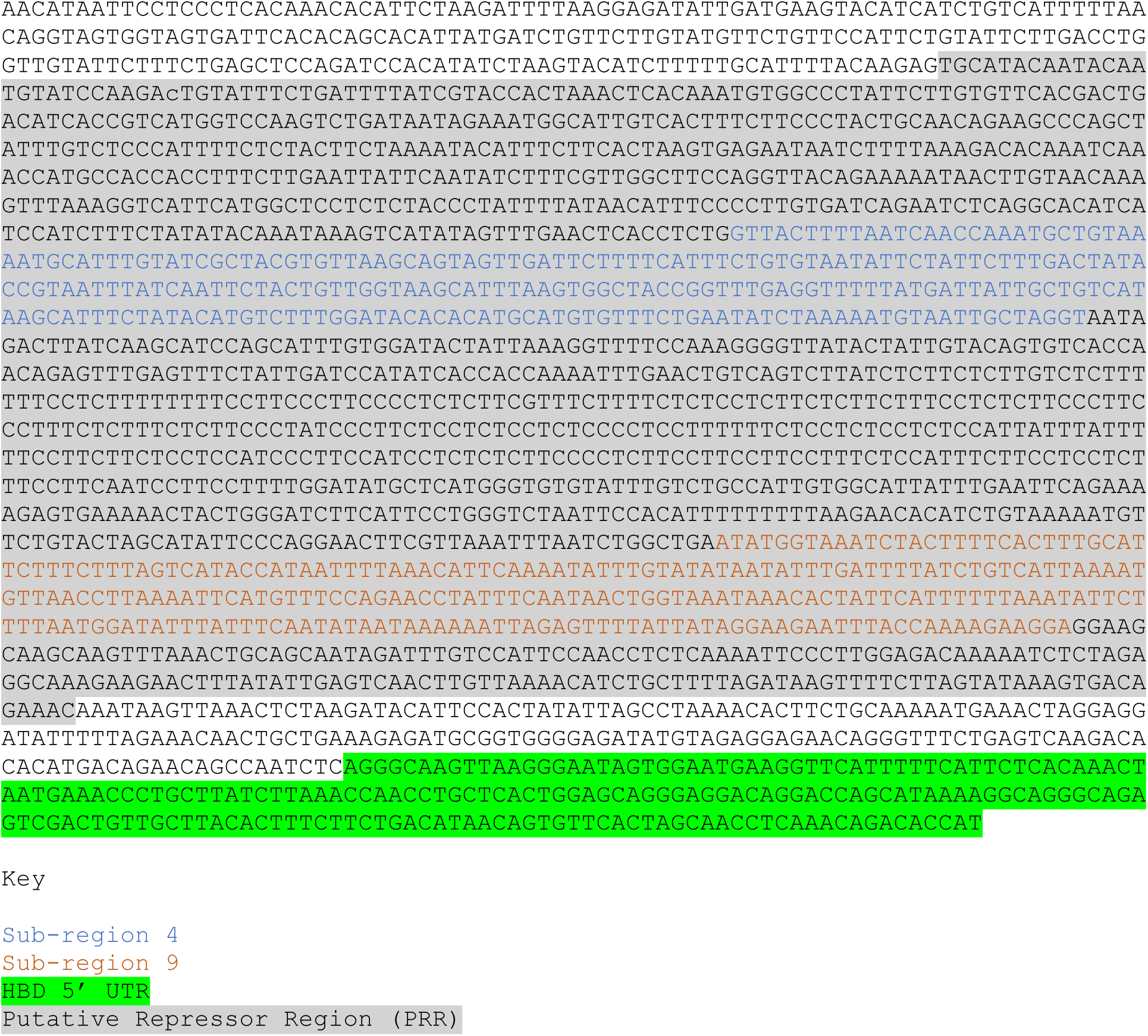

